# Analysis of glycosylation and disulfide bonding of wild-type SARS-CoV-2 spike glycoprotein

**DOI:** 10.1101/2021.04.01.438120

**Authors:** Shijian Zhang, Eden P. Go, Haitao Ding, Saumya Anang, John C. Kappes, Heather Desaire, Joseph Sodroski

## Abstract

The SARS-CoV-2 coronavirus, the etiologic agent of COVID-19, uses its spike (S) glycoprotein anchored in the viral membrane to enter host cells. The S glycoprotein is the major target for neutralizing antibodies elicited by natural infection and by vaccines. Approximately 35% of the SARS-CoV-2 S glycoprotein consists of carbohydrate, which can influence virus infectivity and susceptibility to antibody inhibition. We found that virus-like particles produced by coexpression of SARS-CoV-2 S, M, E and N proteins contained spike glycoproteins that were extensively modified by complex carbohydrates. We used a fucose-selective lectin to enrich the Golgi-resident fraction of a wild-type SARS-CoV-2 S glycoprotein trimer, and determined its glycosylation and disulfide bond profile. Compared with soluble or solubilized S glycoproteins modified to prevent proteolytic cleavage and to retain a prefusion conformation, more of the wild-type S glycoprotein N-linked glycans are processed to complex forms. Even Asn 234, a significant percentage of which is decorated by high-mannose glycans on soluble and virion S trimers, is predominantly modified in the Golgi by processed glycans. Three incompletely occupied sites of O-linked glycosylation were detected. Viruses pseudotyped with natural variants of the serine/threonine residues implicated in O-linked glycosylation were generally infectious and exhibited sensitivity to neutralization by soluble ACE2 and convalescent antisera comparable to that of the wild-type virus. Unlike other natural cysteine variants, a Cys15Phe (C15F) mutant retained partial, but unstable, infectivity. These findings enhance our understanding of the Golgi processing of the native SARS-CoV-2 S glycoprotein carbohydrates and could assist the design of interventions.

## INTRODUCTION

The recently emerged <u>s</u>evere <u>a</u>cute <u>r</u>espiratory <u>s</u>yndrome coronavirus (SARS-CoV-2) is responsible for the ongoing pandemic of COVID-19, a respiratory disease with an estimated 2-5% mortality (1–7). The SARS-CoV-2 spike (S) glycoprotein mediates the entry of the virus into the host cell and influences tissue tropism and pathogenesis (8–13). The S glycoprotein trimer in the viral membrane is the target for neutralizing antibodies, which are important for vaccine-induced protection against infection (9, 11, 12, 14–18). Monoclonal neutralizing antibodies directed against the S glycoprotein are being evaluated as treatments for SARS-CoV-2-infected individuals (15, 19–27). In the virus-producing cell, the S glycoprotein is synthesized in the endoplasmic reticulum, where it assembles into trimers and is initially modified by high-mannose glycans (28, 29). Each of the three SARS-CoV-2 S glycoprotein protomers possesses 22 canonical sequons for N-linked glycosylation (11, 30–36). Coronavirus virions bud into the endoplasmic reticulum-Golgi intermediate compartment (ERGIC), and S glycoprotein trimers on the surface of these virus particles are thought to be processed further during trafficking through the Golgi complex (29, 37–40). In the Golgi, some of the glycans on the S glycoprotein are modified to complex carbohydrates; in addition, the trimeric S glycoprotein is cleaved by furin proteases into S1 and S2 glycoproteins, which associate non-covalently in the virus spike (27–36). During virus entry, the S1 subunit binds the receptor, angiotensin-converting enzyme 2 (ACE2) (9, 11–13, 41–43). The S2 subunit is further processed by host proteases and undergoes extensive conformational changes to mediate the fusion of the viral and target cell membranes (43–47). Following the insertion of the S2 fusion peptide into the host cell membrane, the interaction of two helical heptad repeat regions (HR1 and HR2) on the S2 subunit brings the viral and cell membranes into proximity (44).

The SARS-CoV-2 S glycoprotein trimer is modified by glycosylation, which in other coronaviruses has been suggested to modulate accessibility to neutralizing antibodies as well as host proteases involved in S processing (11, 13, 30–32, 48, 49). Glycans camouflage S glycoprotein peptide epitopes, shielding them from potentially neutralizing antibodies. Glycans can also contribute to epitopes for antibody recognition; for example, the s309 neutralizing antibody interacts with the glycan on Asn 343 of the SARS-CoV-2 S glycoprotein (50).

Virus entry inhibitors and therapeutic or prophylactic neutralizing antibodies must recognize the mature SARS-CoV-2 spike with its natural glycan coat, as it exists on the viral membrane. The glycosylation of the SARS-CoV-2 spike has been studied using soluble or detergent-solubilized versions of the uncleaved S glycoprotein trimer, modified to retain a pretriggered conformation (30, 33–36, 51). Fewer studies of the glycosylation of S glycoproteins on SARS-CoV-2 virion preparations have been conducted (52, 53). Experience with human immunodeficiency virus (HIV-1) indicates that native, membrane-anchored viral envelope glycoproteins can exhibit glycosylation profiles that differ from those of soluble glycoprotein trimers (54–57). Here, we elucidate the glycosylation and disulfide bonding profile of a wild-type SARS-CoV-2 S glycoprotein trimer, and evaluate the importance of naturally occurring variation in O-linked glycans and disulfide bonds. This information enhances our understanding of the complete, functional SARS-CoV-2 S spike and could assist the development and improvement of efficacious therapies, including monoclonal antibodies, and vaccines.

## RESULTS

### Characterization of SARS-CoV-2 S glycoproteins in cell lysates and virus-like particles

We evaluated the wild-type SARS-CoV-2 S glycoprotein expressed alone or in combination with the viral membrane (M), envelope (E) and nucleocapsid (N) proteins, which direct the formation of virus-like particles (VLPs) (58, 59). In the absence of M, E and N proteins, low levels of the S glycoprotein, presumably in extracellular vesicles, were detected in particles prepared by centrifugation of the supernatants of transiently expressing 293T cells (Figure 1A-C). Coexpression of the M, E and N proteins, particularly in combination, resulted in an increase in the level of S glycoprotein in the supernatant pellets. Both uncleaved S and cleaved (S1 and S2) glycoproteins were detected in the particles prepared from the cell supernatants (Figure 1A and B). Two forms of the uncleaved S glycoprotein were detected in the pelleted particles: 1) a faster-migrating form modified by Endoglycosidase Hf (Endo Hf)-sensitive (high-mannose and/or hybrid) glycans, and 2) a more slowly migrating form modified by Endo Hf-resistant (complex) glycans (Figure 1B). Coexpression of the E glycoprotein resulted in an increase in the ratio of complex:high-mannose glycans in the uncleaved S glycoprotein in the pelleted particles. The vast majority of the cleaved S1 and S2 glycoproteins in the pelleted VLPs was resistant to Endo Hf digestion. Thus, in the presence of SARS-CoV-2 M, E and N proteins, the uncleaved and cleaved S glycoproteins on VLPs are largely modified by complex carbohydrates.

**Figure 1.**
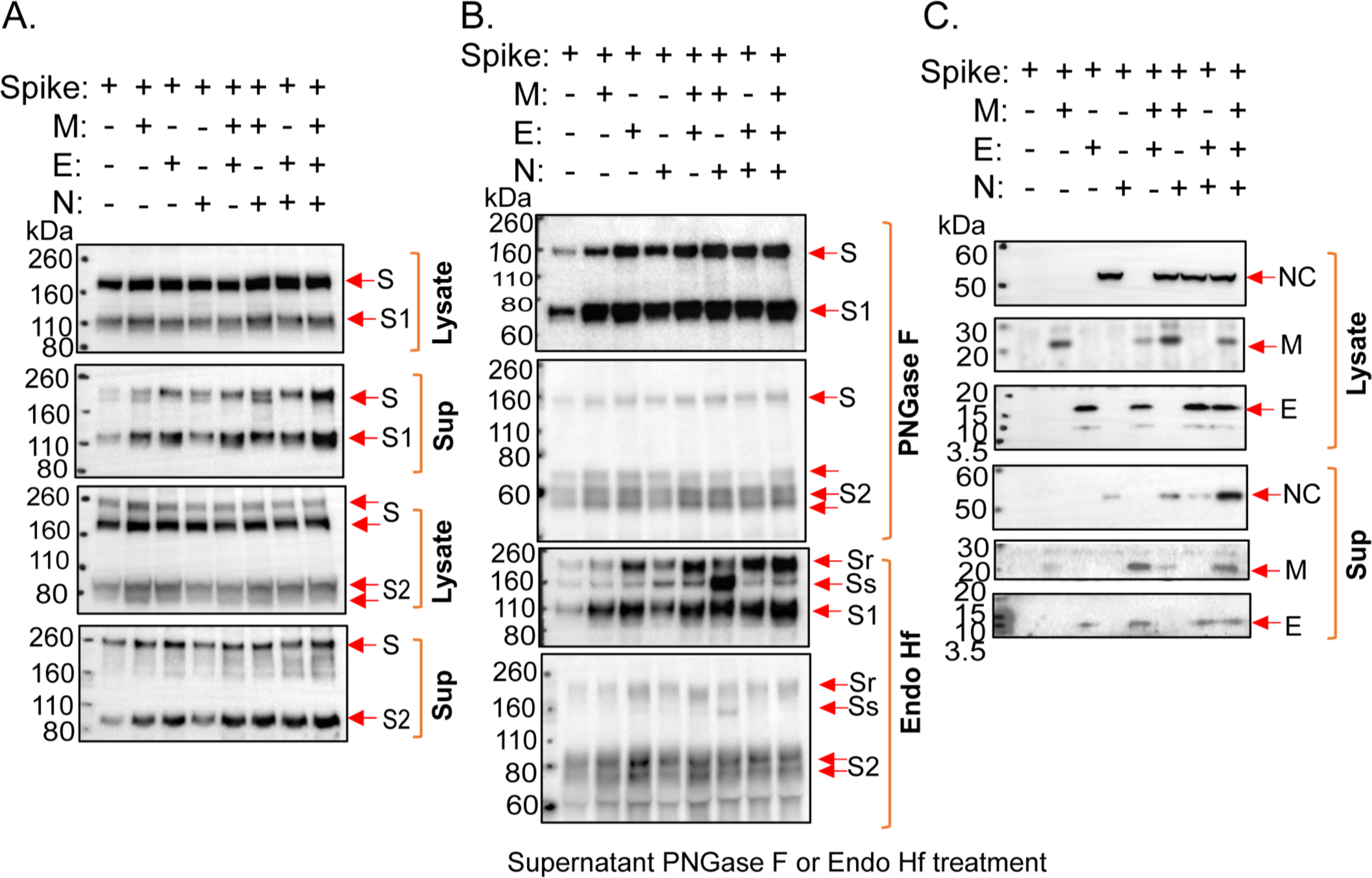
The effect of coexpression of SARS-CoV-2 M, E and N proteins on S glycosylation. (A-C) 293T cells were transfected with plasmids expressing the indicated SARS-CoV-2 proteins (S – spike glycoprotein; M – membrane protein; E – envelope protein; N – nucleocapsid protein). Two days after transfection, cells were lysed and particles were prepared by centrifugation of cell supernatants. (A) Cell lysates and supernatant pellets (Sup) were Western blotted with anti-S1 (upper two panels) and anti-S2 (lower two panels) antibodies. (B) Supernatant pellets were treated with either PNGase F or Endo Hf and then Western blotted with anti-S1 or anti-S2 antibodies. Endo Hf-resistant and Endo Hf-sensitive forms of the S glycoprotein are indicated by S_r_ and S_s_, respectively. (C) Cell lysates and supernatant pellets (Sup) were Western blotted with antibodies against the N, M and E proteins.

### Expression and purification of the SARS-CoV-2 S glycoprotein

To study the native SARS-CoV-2 S glycoprotein in greater detail, we established a stable 293T cell line (293T-S) that expresses the wild-type SARS-CoV-2 S glycoprotein under the control of a tetracycline-inducible promoter (29). To facilitate purification, a carboxy-terminal 2xStrep affinity tag was added to the S glycoprotein, which was otherwise wild-type in sequence (29). Treatment of the 293T-S cells with doxycycline resulted in the expression of the S glycoprotein, which was cleaved into the S1 exterior and S2 transmembrane glycoproteins (Figure 2A). The expressed S glycoproteins mediated the formation of syncytia when human ACE2 (hACE2) was transiently coexpressed in the 293T-S cells (Figure 2A). The expressed S glycoproteins supported infection of 293T-hACE2 cells by a pseudotyped vesicular stomatitis virus (VSV) vector (Figure 2B). Nearly all of the S glycoproteins incorporated into VSV pseudotypes were cleaved (Figure 2B).

**Figure 2.**
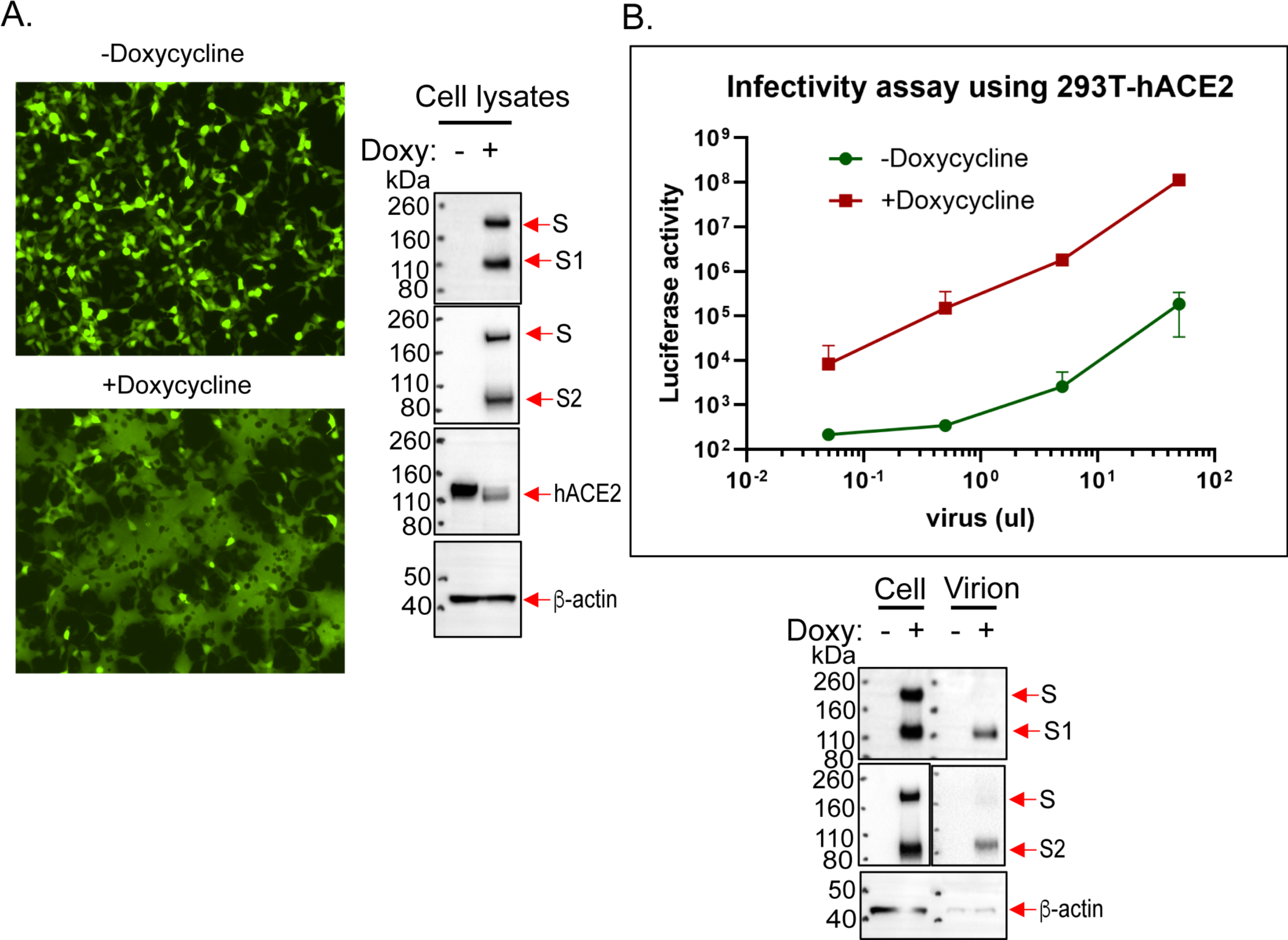
Inducible expression of a functional SARS-CoV-2 S glycoprotein. (A) 293T-S cells were transfected with plasmids expressing eGFP and human ACE2 (hACE2) and then incubated in medium with or without 1 µg/ml doxycycline (Doxy). Two days later, cells were imaged with a fluorescence microscope. Cell lysates were Western blotted with a mouse antibody against S1, a rabbit antibody against S2, a goat antibody against hACE2 and a mouse antibody against α-β-actin. (B) 293T-S cells were cultured for 24 hours in medium with 1 µg/ml doxycycline (Doxy) or in control medium. The cells were then infected with a G glycoprotein-pseudotyped VSVΔG virus encoding luciferase. One day later, virus particles were harvested from pre-cleared cell supernatants and incubated with 293T-hACE2 cells. Luciferase activity was measured one day later. Cell lysates and viruses concentrated by a 100,000 x g centrifugation were Western blotted with a mouse antibody against S1, a rabbit antibody against S2, and an anti-α-β-actin antibody.

For purification of the SARS-CoV-2 S glycoprotein, we evaluated several detergents as well as styrene-maleic acid (SMA) copolymers for their ability to extract the S glycoproteins from 293T-S membranes (60–65). NP-40, Triton X-100 and Cymal-5 solubilized the S glycoproteins more efficiently than lauryl maltyl neopentyl glycol (LMNG) or SMA (Figure 3A). The solubilized S glycoproteins migrated on Blue Native gels at a size consistent with trimers (Figure 3B). Strep-Tactin purification of the cleaved S1/S2 complexes as well as the uncleaved S glycoproteins in Cymal-5 solutions was slightly more efficient than in the other detergents; therefore, we used Cymal-5 to extract the S glycoproteins for purification.

**Figure 3.**
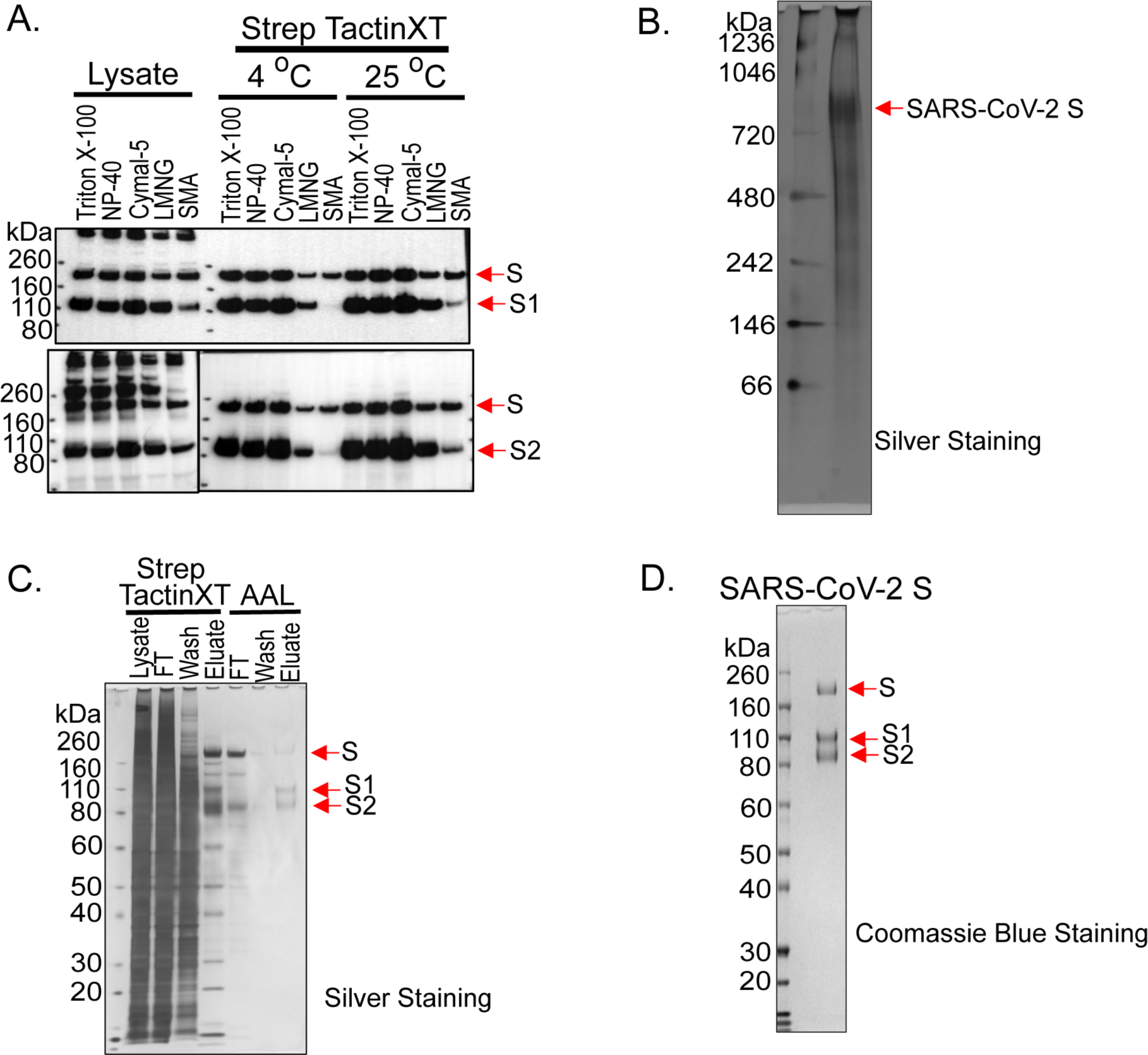
Purification of the SARS-CoV-2 S glycoproteins. (A) 293T-S cells induced with doxycycline for two days were lysed in buffers containing the indicated detergents or styrene-maleic acid (SMA) copolymers. The cell lysates were either directly Western blotted (Lysate) or used for S glycoprotein purification by Strep-Tactin XT at the indicated temperature. The purified S glycoproteins were Western blotted with rabbit antibodies against S1 (upper panel) and S2 (lower panel). (B) Purified S glycoproteins were analyzed on a Blue Native gel, which was stained with silver. (C) A lysate of 293T-S cells in a buffer containing Cymal-5 was purified by Strep-Tactin XT, followed by purification on Aleuria aurantia lectin (AAL)-agarose resin. The samples at various stages of purification were analyzed by SDS-PAGE and silver staining. FT – flow-through fraction. (D) The purified S glycoproteins were analyzed by SDS-PAGE and Coomassie Blue staining.

Both uncleaved and cleaved SARS-CoV-2 S glycoproteins are incorporated into virus-like particles (VLPs) formed as a result of expression of the SARS-CoV-2 M, E and N proteins (59) (Figure 1). The S glycoproteins in VLPs are extensively modified by complex carbohydrates, indicating passage through the Golgi compartment. Therefore, we enriched the mature, Golgi-resident S glycoproteins by sequentially using Strep-Tactin and Aleuria aurantia lectin (AAL) to purify the S glycoproteins from the membranes of 293T-S cells (Figure 3C). AAL recognizes fucose, which is added to a subset of complex glycans in the Golgi apparatus (66–69). The purified S glycoproteins consisted of approximately 25% uncleaved and 75% cleaved (S1 and S2) glycoproteins (Figure 3D).

### Disulfide and glycosylation analysis of the purified S glycoproteins

The disulfide bond topology of the purified S glycoproteins was determined by identifying disulfide-linked peptides from the tryptic digests of the S glycoprotein preparation by mass spectrometry (MS) (Figures 4 and 5). The S1 glycoprotein begins at an N-terminal glutamine (residue 14) that has undergone condensation to form pyroglutamine. The same N-terminus has been observed for secreted, soluble forms of uncleaved SARS-CoV-2 S glycoprotein trimers (33). Ten disulfide bonds in S1 and five disulfide bonds in S2 were mapped. The cysteine residues paired in the mapped disulfide bonds are consistent with those defined by structural analyses (11, 13, 70). We also observed an alternative disulfide bond between Cys 131 and Cys 136 (Figure 5); in current S glycoprotein structures (11, 13, 70), these N-terminal domain cysteine residues are 12-15 Å apart and therefore are unable to form a disulfide bond without a change in conformation. Apparently, a fraction of the expressed S glycoproteins tolerates some plasticity in the N-terminal domain.

**Figure 4.**
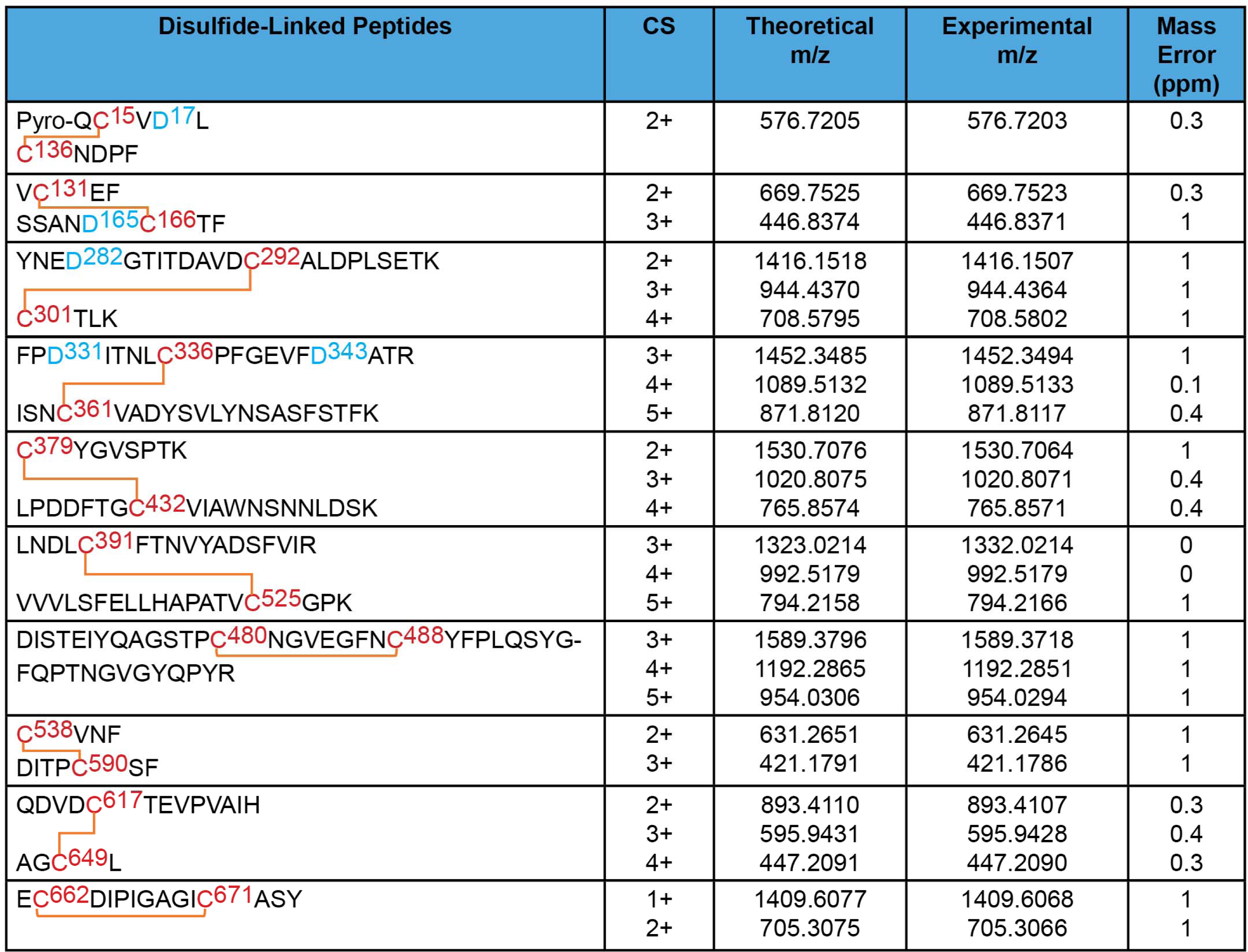
Disulfide bond topology of the purified S1 glycoproteins. MS analysis of the purified S1 glycoproteins identified 10 canonical disulfide bonds between the cysteine residues highlighted in red. Glycosylated asparagine residues converted to aspartic acid residues by PNGase F treatment are highlighted in blue.

**Figure 5.**
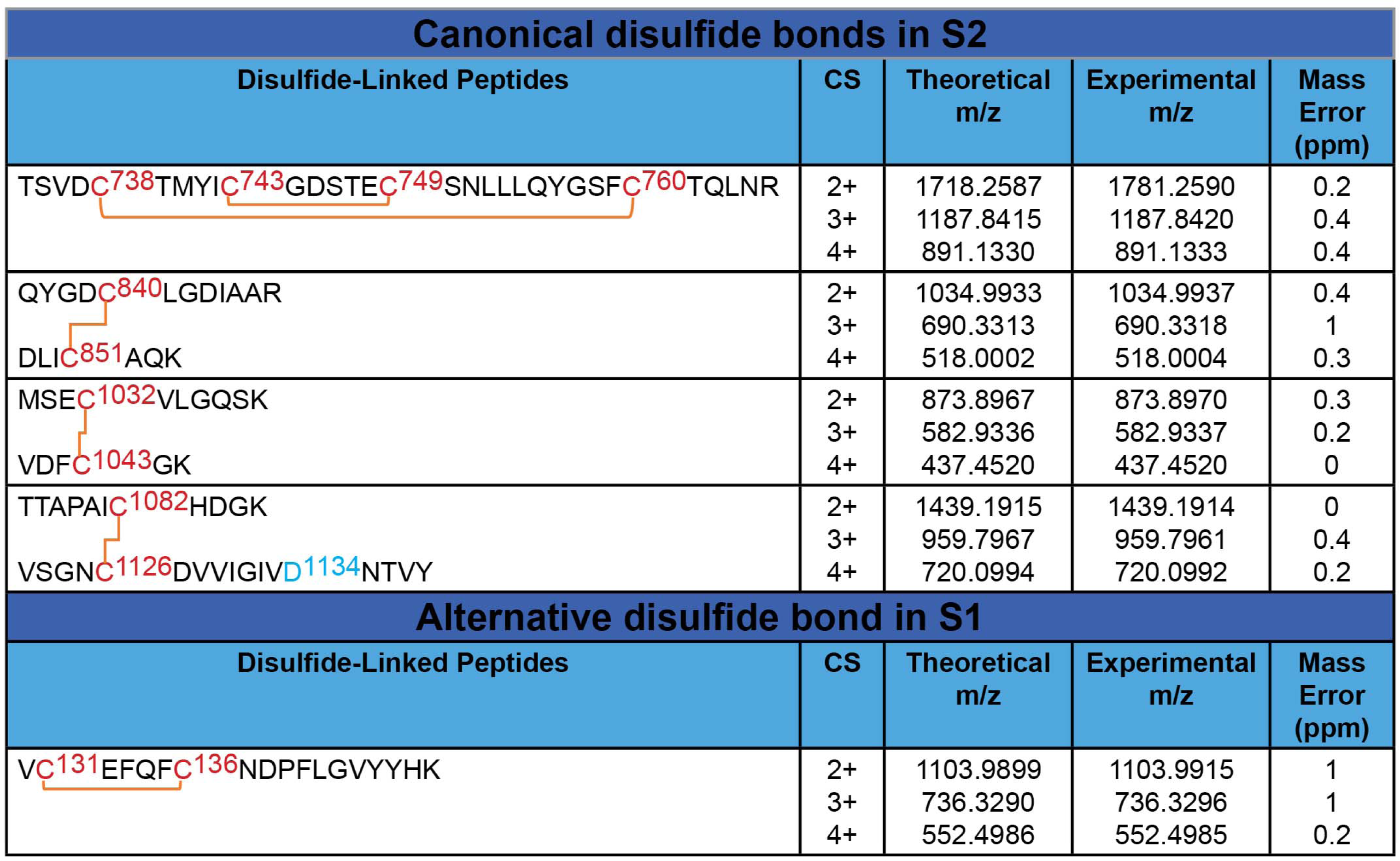
Disulfide bond topology of the purified S glycoproteins. MS analysis of the purified S2 glycoproteins identified 5 canonical disulfide bonds (upper panel). The MS analysis also identified one alternative disulfide bond in the S1 glycoprotein (lower panel). Cysteine residues participating in the disulfide bonds are highlighted in red. The glycosylated asparagine residue converted to an aspartic acid residue by PNGase F treatment is highlighted in blue.

The glycan profile and glycosylation site occupancy of the 22 potential N-linked glycosylation sites were determined using an integrated glycopeptide-based MS approach described previously (54, 55, 71). With the exception of one site, Asn 1074, all of the N-linked glycosylation sites on this protein were fully occupied with glycans (Table 1). Asn 1074 was detected as partially occupied, although the unoccupied form is just one of over fifty different forms present at this site.

**Table 1.** Glycosylation site occupancy. The sequences of the SARS-CoV-2 glycopeptides with N-linked glycosylation (A) and O-linked glycosylation (B) are shown, with the potential sites of glycosylation highlighted in red and green, respectively. The occupancy at each potential N-linked glycosylation (PNG) or O-linked glycosylation site is indicated.

|  | No of PNG Sites | Occupancy |
| --- | --- | --- |
| Pyro_QCVN<sup>17</sup>LTTR | 1 | 1 |
| FSN<sup>61</sup>VTWF | 1 | 1 |
| HAIHVSGTN<sup>74</sup>GTK | 1 | 1 |
| TQSLLIVN<sup>122</sup>ATNVVIK/IVNN<sup>122</sup>ATNVVIK | 1 | 1 |
| HKN<sup>149</sup>NKSWMESEF/N<sup>149</sup>NK | 1 | 1 |
| VYSSANN<sup>165</sup>CTFEYVSQPFLMDLEGK/SSANN<sup>165</sup>CTF | 1 | 1 |
| DLPQGFSALEPLVDLPIGIN<sup>234</sup>ITR/SALEPLVDLPIGIN<sup>234</sup>ITR | 1 | 1 |
| YNEN<sup>282</sup>GTITDAVDCALDPLSETK | 1 | 1 |
| FPN<sup>331</sup>ITNLCPF/FPN<sup>331</sup>ITNL | 1 | 1 |
| GEVFN<sup>343</sup>ATRF/ N<sup>343</sup>ATRF | 1 | 1 |
| GGVSVITPGTN<sup>603</sup>TSNQVAVLY | 1 | 1 |
| QDVN<sup>616</sup>CTEVPVAIHADQLTPTWR/ QDVN<sup>616</sup>CTEVPVAIHADQL | 1 | 1 |
| AGCLIGAEHVN<sup>657</sup>NSYECDIPIGAGICASYQTQTNSPR/AGCLIGAEHVN<sup>657</sup>NSY | 1 | 1 |
| SVASQSIIAYTMSLGAENSVAYS<sup>N709</sup>NSIAIPTN<sup>717</sup>FTISVTTEILPVSMTK/S<sup>N709</sup>NSIAIPTN<sup>717</sup>F | 2 | 2 |
| DFGGFN<sup>801</sup>FSQILPDPSKPSK/TPPIKDFGGFN<sup>801</sup>FSQILPDPSKPSK | 1 | 1 |
| GYHLMSFPQSAPHGVVFLHVTYVPAQEK<sup>N1074</sup>FTTAPAICHDGK / N<sup>1074</sup>FTTAPAICHDGK / VPAQEK<sup>N1074</sup>F | 1 | 0 and 1 |
| EGVFVSN<sup>1098</sup>GTHWFVTQR/VFVSN<sup>1098</sup>GTHW | 1 | 1 |
| VSGNCDVVIGIVN<sup>1134</sup>NTVY | 1 | 1 |
| N<sup>1158</sup>HTSPDVLGDISGIN<sup>1173</sup>ASVVNIQK | 2 | 2 |
| NLN<sup>1194</sup>ESLIDLQELGK | 1 | 1 |

**B. O-Linked Glycosylation**
|  | No of O-linked Sites | Occupancy |
| --- | --- | --- |
| VQPT<sup>323</sup>ES<sup>325</sup>IVR | 2 | 0 and 1 |
| AGCLIGAEHVNNS<sup>659</sup>YECDIPIGAGICAS<sup>673</sup>YQT<sup>676</sup>Q T<sup>678</sup>NS<sup>680</sup>PR | 5 | 0,1 and 2 |
| T<sup>696</sup>MSLGAENSVAY | 1 | 0 and 1 |
| NHT<sup>1160</sup>SPDVDLGDIS<sup>1170</sup>GINASVVNIQK | 2 | 0 and 1 |

A pictorial description of the glycosylation profile of this protein is shown in Figure 6. In sum, 826 unique N-linked glycopeptides were detected, along with 17 O-linked glycopeptides. This glycosylation coverage is more in-depth than early reports, where the number of unique glycoforms detected was more typically in the 100-200 range (30, 33, 35). Furthermore, this analysis provides the first report of O-linked glycosylation at 7 glycosylation sites: S659, S673, T676, S680, T696, T1160,and S1170.

**Figure 6.**
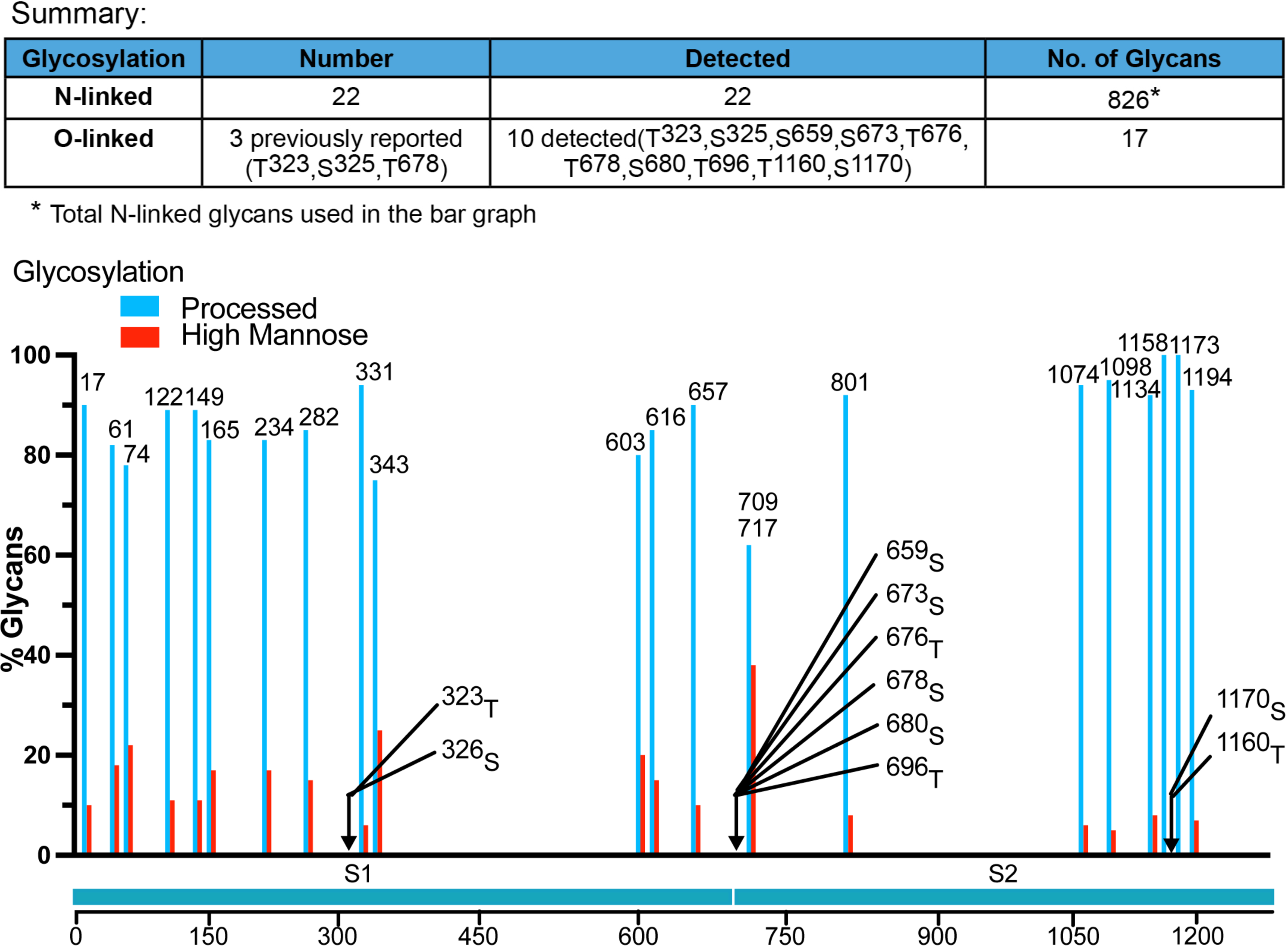
Glycosylation profile of the purified S glycoproteins. MS analysis of the purified S glycoproteins identified 22 N-linked glycosylation sites as well as O-linked glycosylation sites, summarized in the upper panel. The glycan composition at each N-linked glycosylation site is shown in the lower panel. Serine and threonine residues contained in glycopeptides with O-linked carbohydrates are indicated by arrows.

As can be seen in Figure 6, processed glycoforms predominated at each N-linked glycosylation site and, as shown in Figure 7A, these complex forms were highly, but not exclusively, fucosylated. Additionally, even though each glycosylation site was heavily processed in the Golgi, the sialic acid content varied across the protein sequence. Some sites, like N61 and N603 had no sialylated glycoforms detected, while most of the sites in the S2 protein, particularly those nearest the C terminus, were abundantly sialylated. Finally, as shown in Figure 7B, a number of O-linked glycoforms were detected.

**Figure 7.**
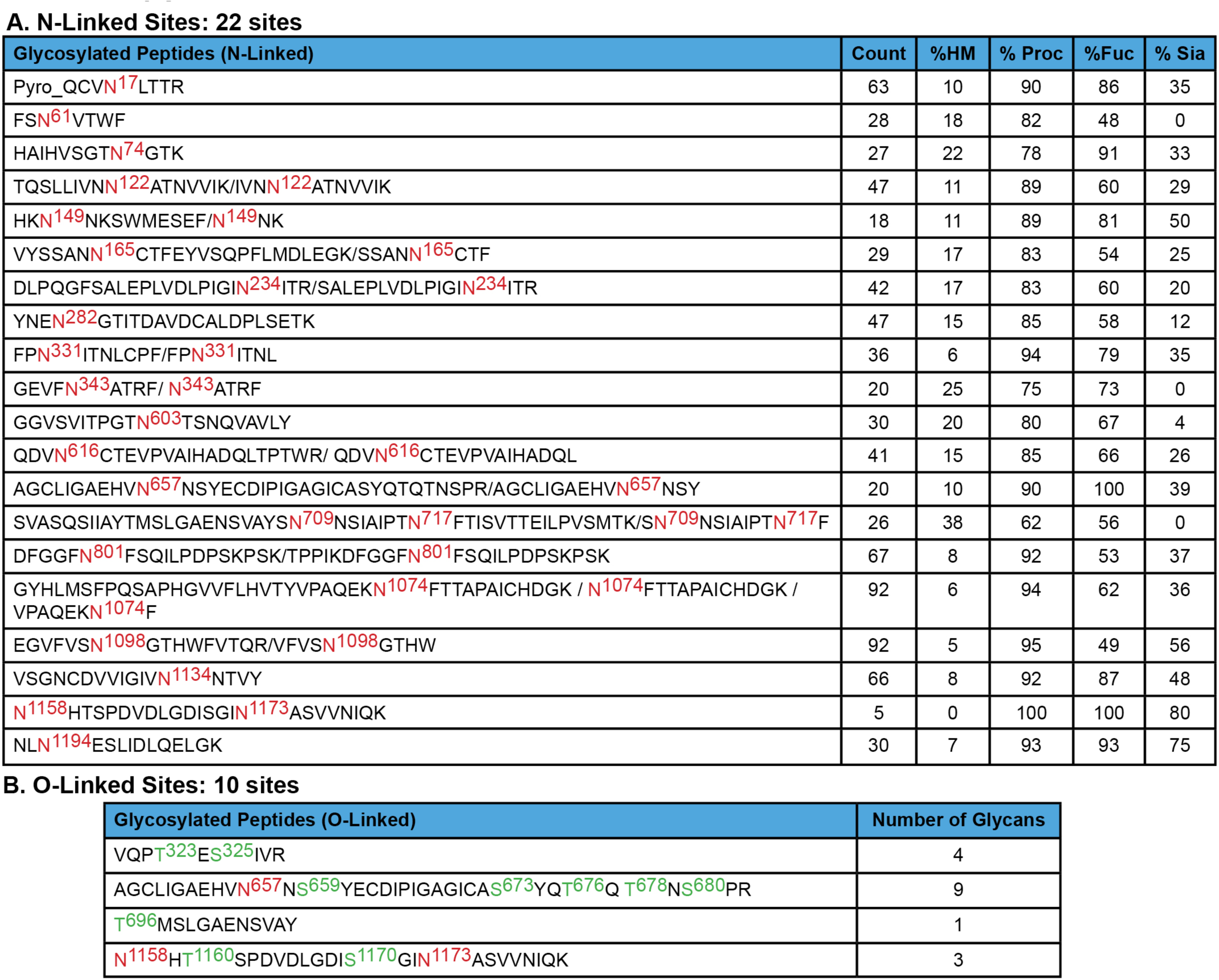
SARS-CoV-2 S glycopeptides. (A) The sequences of the S glycopeptides with N-linked glycosylation are shown, with the modified asparagine residues highlighted in red. The percentage of glycans that are high-mannose (HM), processed (Proc) (complex + hybrid), modified by fucose (Fuc) or sialylated (Sia) are indicated. (B) The sequences of S glycopeptides with O-linked glycosylation are shown, with the potentially modified serine and threonine residues highlighted in green. Asparagine residues in the glycopeptides that are modified by N-linked glycans are highlighted in red.

The vast majority of the N-linked glycans on the Golgi-enriched, wild-type SARS-CoV-2 S glycoprotein expressed in 293T cells were processed to complex glycans. In Figure 8, we compare our results with available glycosylation analyses of soluble or solubilized SARS-CoV-2 S glycoproteins. These include wild-type S glycoproteins purified from SARS-CoV-2 virions, as well as soluble and full-length S glycoprotein trimers modified to inhibit furin cleavage and to stabilize a prefusion conformation (30, 33, 34, 36, 51–53). The glycans on individually expressed SARS-CoV-2 S1 and S2 glycoproteins have also been analyzed (35). All studies agree that complex carbohydrates are found on most of the N-linked glycan sites on SARS-CoV-2 S glycoprotein trimers, as well as on recombinant soluble S1 and S2 glycoproteins. However, the extent of N-linked glycan processing in our study is greater than that seen for either soluble S glycoprotein trimers or for modified (cleavage-defective, proline-substituted) S glycoprotein variants (30, 33, 34, 36, 51). The glycosylation profile of our Golgi-enriched S glycoprotein preparation most closely resembles that of S glycoproteins purified from SARS-CoV-2 virions propagated in Vero cells (Figure 8). However, in our S glycoprotein preparation, Asn 234 in the S1 N-terminal domain is mostly processed, whereas high-mannose glycans are retained at this site in the other characterized S glycoprotein variants. The highest composition of high-mannose glycans in our study mapped to a glycopeptide containing two potential N-linked glycosylation sites at Asn 709 and Asn 717. Although we cannot precisely assign a glycan composition at either site, our result is consistent with observations on soluble/modified S glycoproteins suggesting that one or both of these sites is occupied by a significant percentage of high-mannose glycans (30, 33, 34). The location of these glycans in a heavily glycosylated region near the base of the S1 subunit may limit the access of glycosylation enzymes and predispose to the retention of high-mannose glycans (Figure 9).

**Figure 8.**
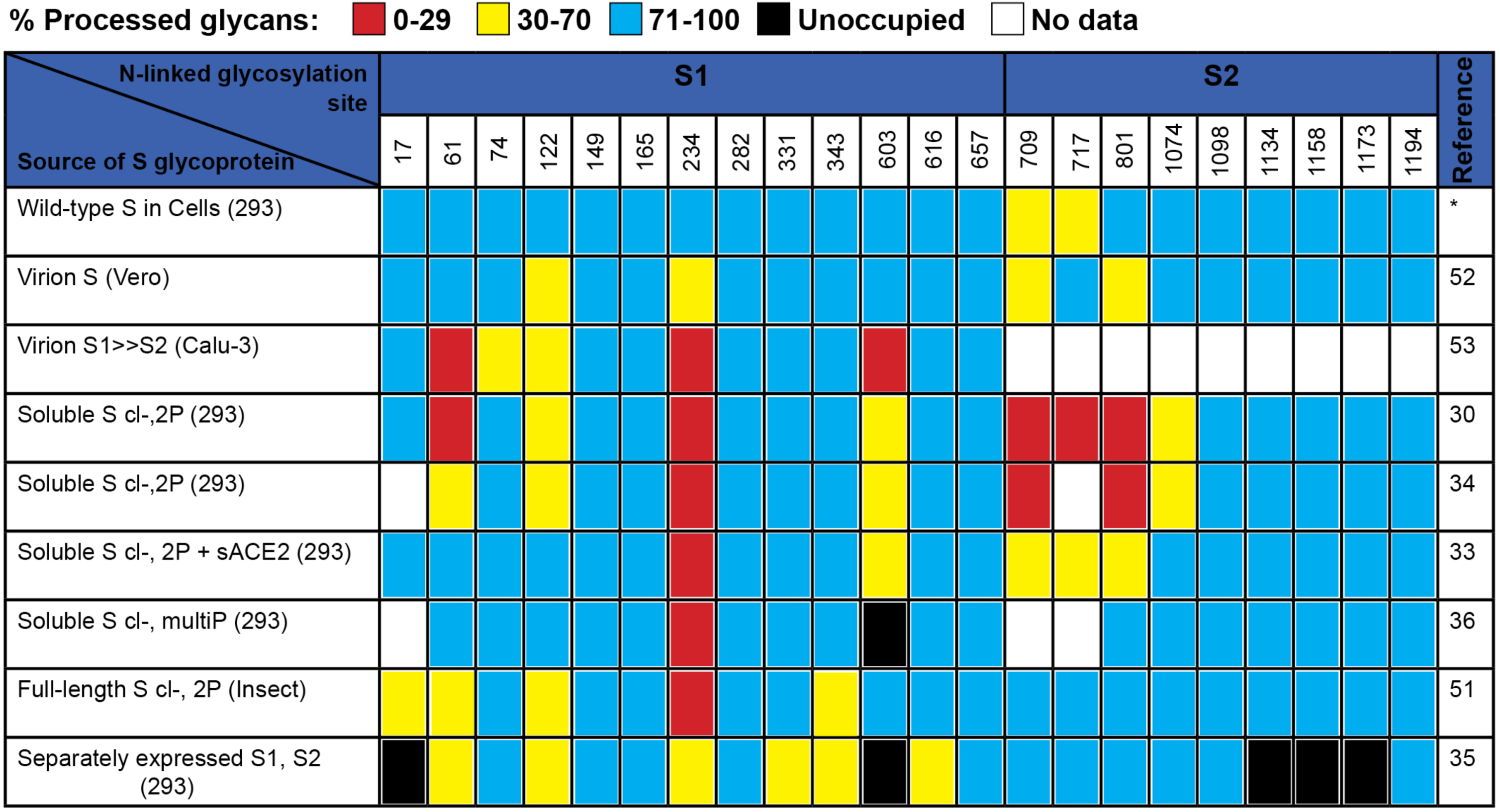
Glycan composition of different SARS-CoV-2 S glycoprotein preparations. The glycan composition at each of the potential S glycoprotein N-linked glycosylation sites from this study (upper row) (∗) is compared with those defined for SARS-CoV-2 S glycoproteins from various sources. Some of the S glycoproteins have been produced in soluble forms with alterations of the furin cleavage site (cl-) and with proline substitutions (2P or multiP) to stabilize pre-fusion conformations.

**Figure 9.**
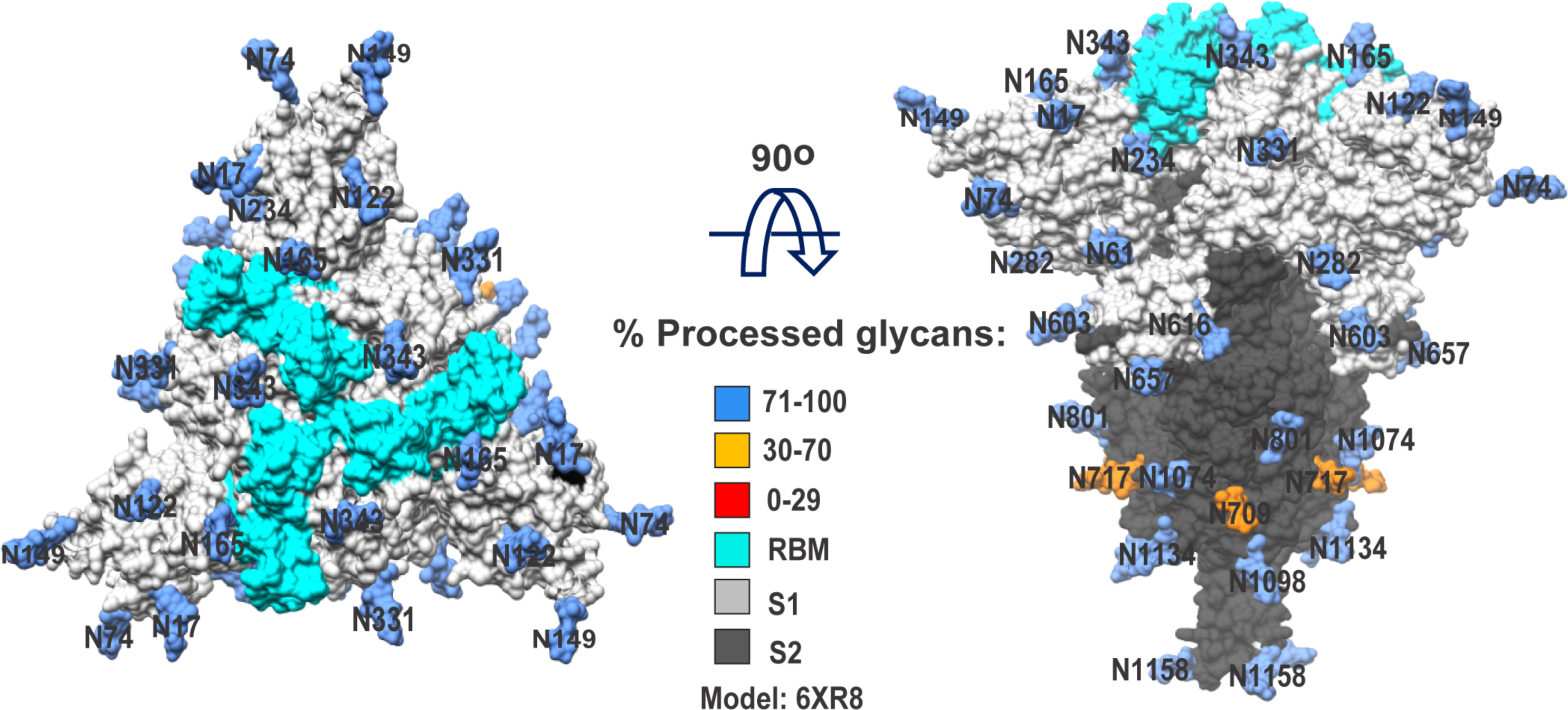
Location of glycans on the SARS-CoV-2 S glycoprotein trimer. N-linked glycans associated with the indicated asparagine residues are shown on the cryo-EM structure of a solubilized SARS-CoV-2 S glycoprotein trimer (PDB 6XR8) (70). The S1 subunits are colored light grey, and the S2 subunits are colored dark grey. The receptor-binding motif is colored cyan. The glycans are colored according to the level of processing observed in our study.

O-linked glycosylation of two S1 glycopeptides and two S2 glycopeptides was detected (Figure 7B). Potential candidates for O-glycosylated residues include Thr 323, Ser 325, Ser 659, Ser 673, Thr 676, Thr 678 and Ser 680 in S1, and Thr 696, Thr 1160 and Ser 1170 in S2. O-linked glycosylation at Thr 323/Ser 325 was reported for soluble S glycoproteins and virion S glycoprotein trimers, but with low occupancy; in some cases, less than 1% of the residues were modified (33, 35, 36, 53). In this study, the occupancy rate for these two sites is also around 1% (Table 1). Thr 678 has also been reported to be O-glycosylated in soluble and virion S glycoproteins, with higher occupancy than that for Thr 323/Ser 325 (36, 53). In our study, the peptide containing Ser 659, Ser 673, Thr 676, Thr 678 and Ser 680 was found to be occupied by at least one O-linked glycan at a level of about 5%.

### SARS-CoV-2 S glycoprotein expression in a cell line with defective glycosylation

GALE/GALK2 293T cells defective for O-linked glycosylation have been established (73). Both the wild-type SARS-CoV-2 S glycoprotein and the prevalent D614G variant S glycoprotein (29, 74–76) were expressed in 293T cells and in the GALE/GALK2 293T cells. Both the S1 and S2 glycoproteins of the wild-type and D614G SARS-CoV-2 strains migrated faster when expressed in the GALE/GALK2 293T cells compared with the migration of these glycoproteins expressed in 293T cells (Figure 10A). Digestion of the glycoproteins with PNGase F revealed that the observed differences in S1 migration could be explained by differences in N-linked glycosylation. PNGase F digestion of the S2 glycoprotein resulted in 60- and 63-kD products. The 60-kD PNGase F-produced S2 proteins expressed in wild-type and GALE/GALK2 293T cells migrated similarly, ruling out a significant level of O-linked glycosylation. The 63-kD PNGase F S2 product potentially is O-glycosylated, as it migrated faster when synthesized in the GALE/GALK2 293T cells. However, the 63-kD product observed after PNGase F digestion was minimally affected by further treatment with O-glycosidase + neuraminidase. Moreover, no significant difference in the migration of the untreated S1 and S2 glycoproteins was observed after O-glycosidase + neuraminidase treatment. We note that the Core 1 O-linked glycans detected on the purified S glycoproteins should be digestible by O-glycosidase + neuraminidase, as was shown on a control substrate, fetuin (Figure 10B). Taken together, these results indicate that O-glycan occupancy is low on the SARS-CoV-2 S glycoproteins expressed in 293T cells. Variation in other post-translational modifications, including N-linked glycosylation, apparently accounts for most of the observed differences in the migration of S1 and S2 glycoproteins expressed in wild-type and GALE/GALK2 293T cells. When the S glycoproteins produced in GALE/GALK2 293T cells were used to pseudotype vesicular stomatitis virus (VSV) vectors, the resulting viruses exhibited lower infectivity than viruses made with S glycoproteins produced in 293T cells (data not shown). These results suggest that differences in the post-translational modifications of S glycoproteins produced in 293T and GALE/GALK2 293T cells may influence S glycoprotein function. As only a fraction of the S glycoprotein is modified by O-linked glycans in 293T cells, differences in O-linked glycosylation are unlikely to explain the observed reduction in the infectivity of VSV(S) pseudotypes produced in GALE/GALK2 293T cells.

**Figure 10.**
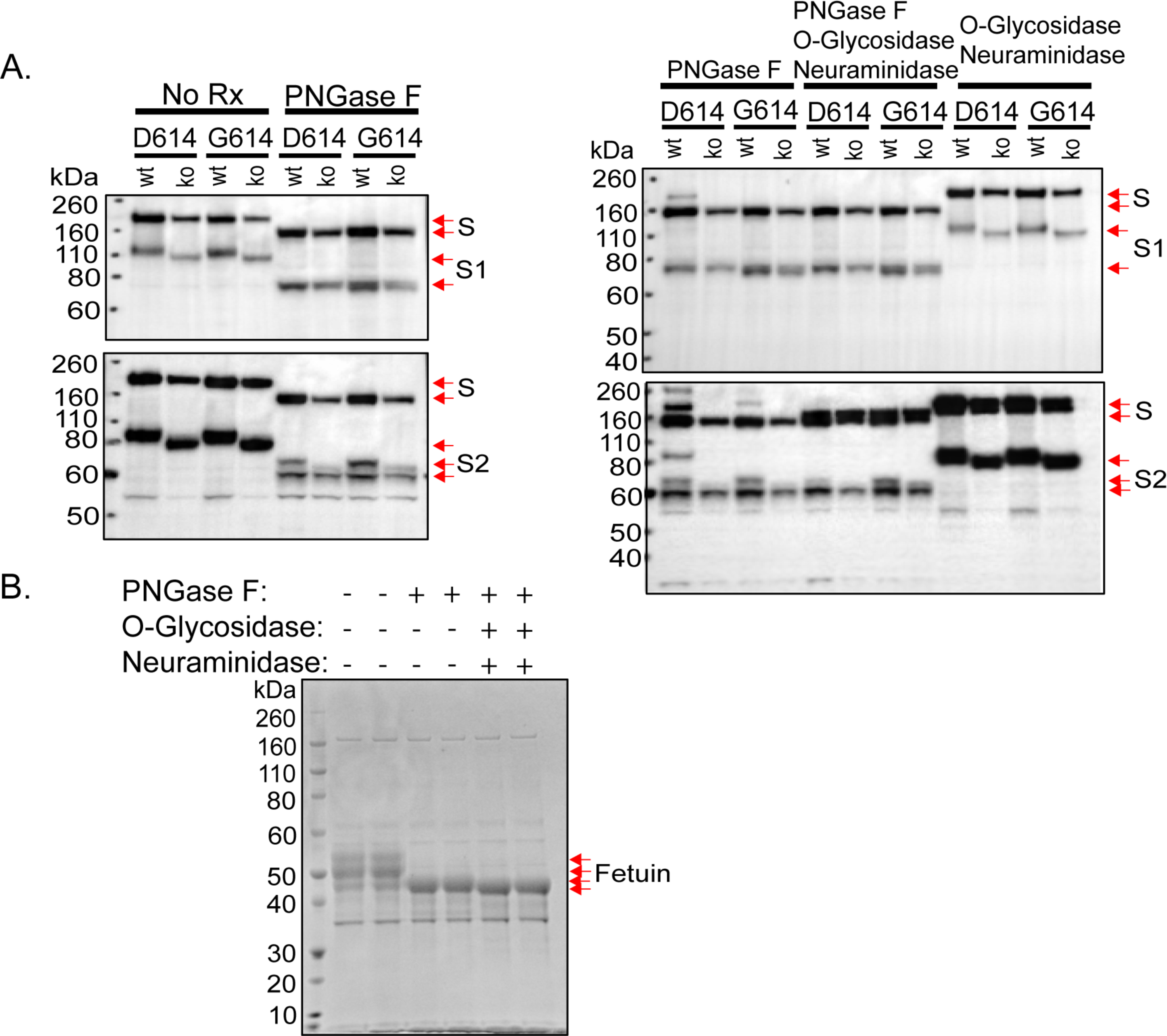
Characterization of wild-type and D614G S glycoproteins in GALE/GALK2 293T cells. (A) The wild-type SARS-CoV-2 S glycoprotein (D614, with an aspartic acid residue at 614) and the D614G variant (G614, with a glycine residue at 614) were expressed in wild-type 293T cells (wt) or in GALE/GALK2 293T cells (ko) (73). Cell lysates were untreated (No Rx) or were treated with the indicated glycosidase(s), followed by Western blotting with a mouse antibody against S1 (upper panels) or a rabbit antibody against S2 (lower panels). The S glycoproteins, either untreated or treated with different glycosidases, are indicated by red arrows. (B) As a control, fetuin, which has N- and O-linked glycans, was treated with the indicated glycosidases. The different fetuin glycoforms are indicated by red arrows.

### Natural variants of SARS-CoV-2 S glycoproteins

Rare natural variants of SARS-CoV-2 S glycoproteins exhibit substitutions at one of the cysteine residues, potentially compromising the formation of particular disulfide bonds (77, 78). We wished to assess the impact of these substitutions on S glycoprotein expression, processing and function. The C15F change eliminates the Cys 15-Cys 136 disulfide bond in the S1 N-terminal domain. Despite this alteration, the C15F S glycoprotein was proteolytically processed nearly as efficiently as the wild-type S glycoprotein and exhibited wild-type association of the S1 and S2 subunits (Figure 11 and Table 2). The infectivity of VSV vectors pseudotyped with the C15F S glycoproteins was approximately 31% of that of virus pseudotyped with the wild-type S glycoproteins. However, after freeze-thawing, the relative infectivity of the C15F mutant virus decreased dramatically (data not shown). Apparently, the Cys 15-Cys 136 disulfide bond is not absolutely essential for S glycoprotein function, but may contribute to the stability of the functional spike. By contrast, the C301F and C379F changes, which eliminate the Cys 291-Cys 301 and Cys 379-Cys 432 disulfide bonds respectively located in the S1 N-terminal domain and receptor-binding domain, resulted in S glycoproteins that were not processed into S1 and S2 glycoproteins (Figure 11 and Table 2). Viruses pseudotyped with the C301F and C379F S glycoproteins exhibited very low levels of infectivity. Thus, of these rare cysteine variants of the SARS-CoV-2 S glycoprotein, only one (C15F) allows partial, but unstable, infectivity.

**Figure 11.**
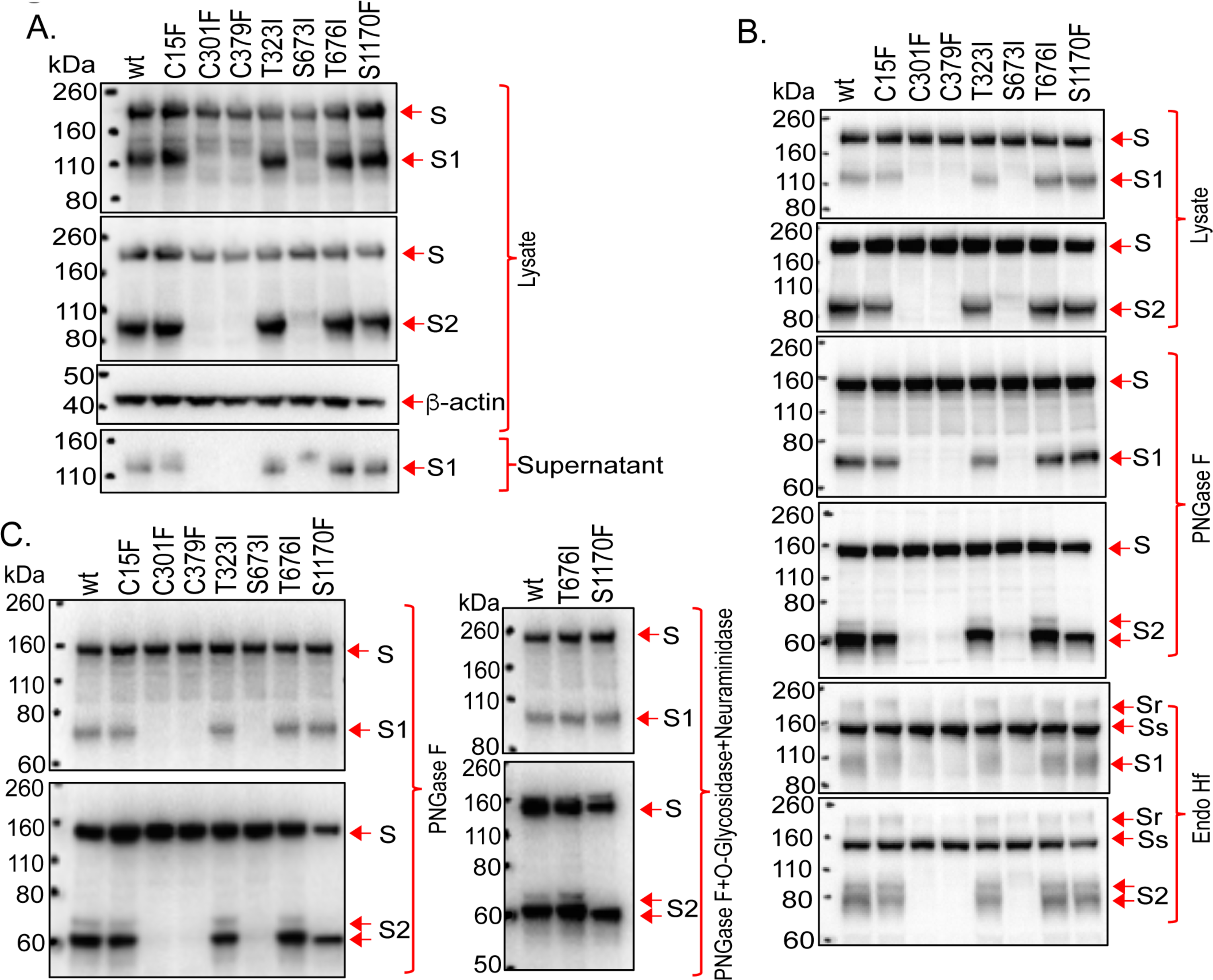
Phenotypes of natural S glycoprotein variants. (A-C) The wild-type SARS-CoV-2 S glycoprotein and the indicated mutants were expressed in 293T cells. Cell lysates were prepared and, in some cases, treated with PNGase F, Endo Hf or PNGase F + O-glycosidase + neuraminidase. The cell lysates were Western blotted. In A, cell supernatants were also collected, precleared by centrifugation at 1800 x g for 10 minutes and used for precipitation by a 1:100 dilution of NYP01 convalescent serum and Protein A-agarose beads. The processing and subunit association indices shown in Table 1 were calculated for each mutant and were normalized to those of the wild-type (wt) S glycoprotein. In B and C, the effects of glycosidases on the wild-type and mutant S glycoproteins in cell lysates are shown. Endo Hf-resistant (S_r_) and -sensitive (S_s_) forms of the uncleaved S glycoprotein are indicated.

**Table 2.**
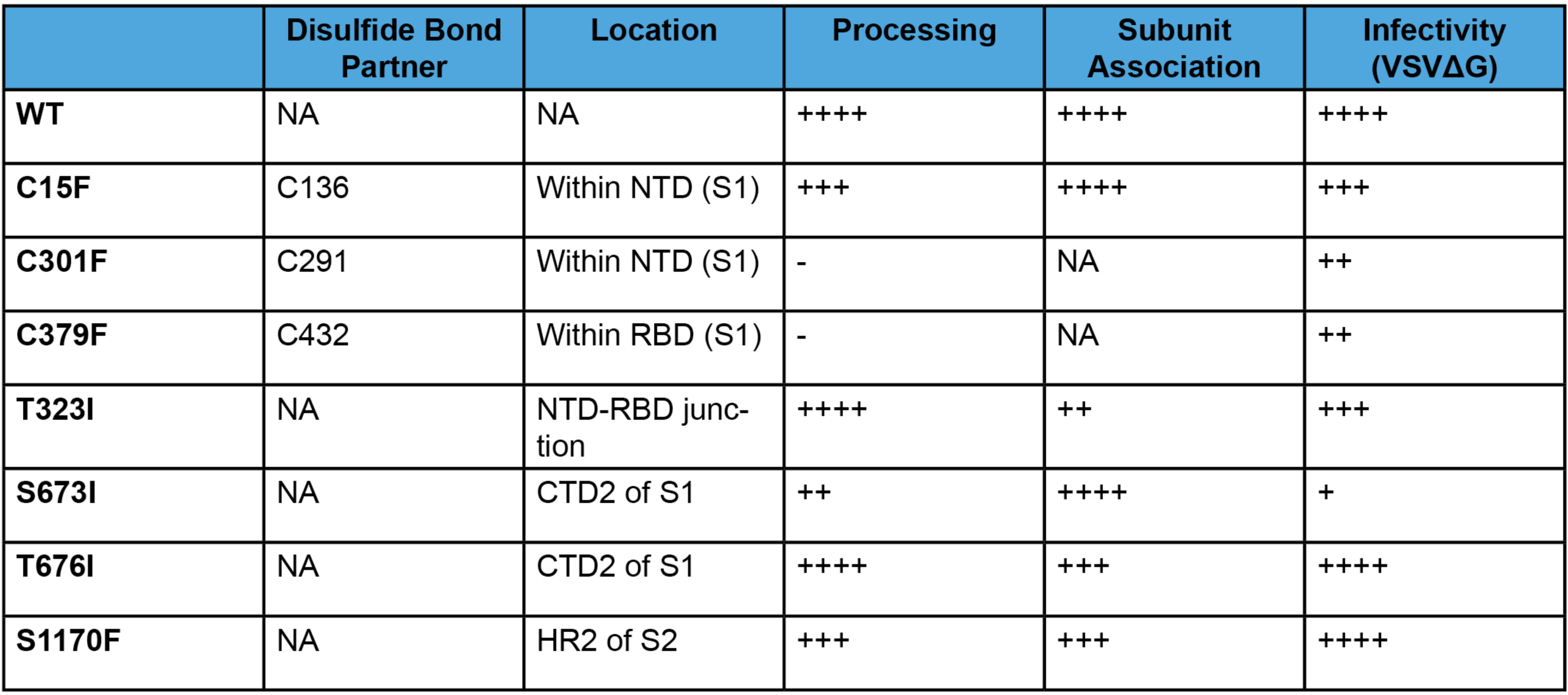
Phenotypes of natural S glycoprotein variants. The processing, subunit association and infectivity of the mutant S glycoproteins, relative to those of the wild-type S glycoprotein, are shown (–, undetectable; +, 1-10% of wild-type level; ++, 11-30% of wild-type level; +++, 31-80% of wild-type level; ++++, 81-120% of wild-type level; NA – not applicable). The location of the altered amino acid residue in the S glycoprotein is indicated: NTD – N-terminal domain; RBD – receptor-binding domain; CTD2 – C-terminal domain 2; HR2 – heptad repeat 2.

|  | Disulfide Bond Partner | Location | Processing | Subunit Association | Infectivity (VSVΔG) |
| --- | --- | --- | --- | --- | --- |
| WT | NA | NA | ++++ | ++++ | ++++ |
| C15F | C136 | Within NTD (S1) | +++ | ++++ | +++ |
| C301F | C291 | Within NTD (S1) | - | NA | ++ |
| C379F | C432 | Within RBD (S1) | - | NA | ++ |
| T323I | NA | NTD-RBD junction | ++++ | ++ | +++ |
| S673I | NA | CTD2 of S1 | ++ | ++++ | + |
| T676I | NA | CTD2 of S1 | ++++ | +++ | ++++ |
| S1170F | NA | HR2 of S2 | +++ | +++ | ++++ |

To evaluate the potential O-linked glycosylation of the SARS-CoV-2 S glycoprotein, several of the threonine and serine residues implicated in our MS study were altered to residues found in natural SARS-CoV-2 variants (77, 78). The T323I, S676I and S1170F mutants were processed nearly as efficiently as the wild-type S glycoprotein, and exhibited good subunit association (Figure 11A and Table 2). The 63-kD S2 glycoprotein band seen in PNGase F-treated lysates from 293T cells expressing the wild-type S glycoproteins was not evident in lysates from cells expressing the S1170F mutant (Figure 11B and C). As the S1170F change does not alter a potential N-linked glycosylation site, it apparently affects other post-translational modifications; as discussed above, resistance of the 63-kD PNGase F product to O-glycosidase appears to rule out modification by Core 1 or Core 3 O-glycans (Figure 11C). The S676I and S1170F mutants supported the entry of VSV pseudotypes as efficiently as the wild-type S glycoprotein (Table 2). The T323I-pseudotyped viruses infected cells with approximately 41% of the efficiency of viruses pseudotyped with the wild-type S glycoproteins, but the infectivity of these viruses decreased further upon freeze-thawing. The S673I mutant was processed inefficiently and only supported the infection of pseudotyped VSV vectors at a very low level.

We examined the sensitivity of the two most replication-competent S glycoprotein mutants, T676I and S1170F, to neutralization by soluble ACE2 (sACE2) and sera from convalescing SARS-CoV-2-infected individuals. No significant differences in the neutralization sensitivity of the wild-type and mutant viruses were observed (Figure 12).

**Figure 12.**
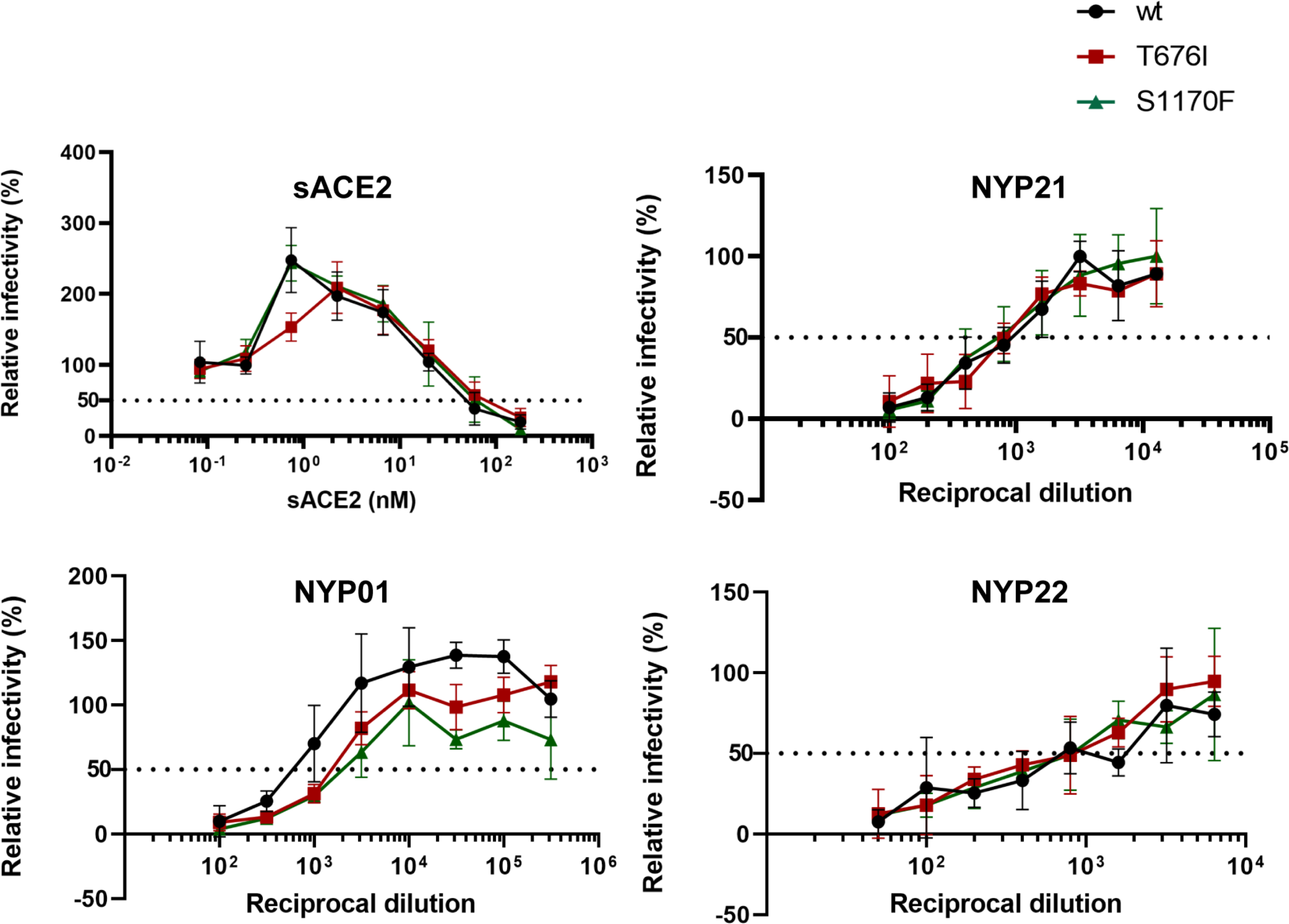
Neutralization of viruses pseudotyped with S glycoprotein variants. The sensitivity of VSV pseudotypes with the indicated S glycoproteins to neutralization by sACE2 or the NYP01, NYP21 and NYP22 convalescent-phase sera is shown. The infectivity is shown relative to that of a mock-treated virus.

## DISCUSSION

As the extensive glycosylation of the spike (S) glycoprotein can potentially influence SARS-CoV-2 infectivity and sensitivity to antibody inhibition, an understanding of the glycosylation profile of the native S glycoprotein trimer is valuable. Glycosylation of the SARS-CoV-2 S glycoprotein can apparently be influenced by subcellular localization and the coexpression of viral proteins. Because proteolytic activation of the S glycoprotein can occur at the cell surface or in endosomal compartments during virus entry, the uncleaved S glycoprotein, as well as the cleaved S glycoproteins, on virions can support virus infection (29, 45–47). Both uncleaved and cleaved S glycoproteins on virions are extensively modified by complex carbohydrates, indicating passage through the Golgi complex (29, 69, 79). We found that coexpression of the SARS-CoV-2 E protein led to a further enrichment of complex glycans on the uncleaved S glycoprotein on VLPs. These observations provided a rationale for focusing on the Golgi-resident S glycoproteins. By including a lectin, AAL, that recognizes fucose in the purification scheme, we attempted to increase the representation of S glycoproteins that passed through the Golgi complex, where fucosylation occurs (66–69). This purification strategy allowed an evaluation of the glycan composition of a Golgi-enriched subset of the S glycoproteins synthesized in the expressing cell; it is conceivable that less processed forms of the S glycoproteins might also be present on virions, depending on host cell type, production levels and VLP characteristics.

SARS-CoV-2 S glycoproteins produced for use as vaccine immunogens have been designed to allow secretion of soluble trimers, to inhibit furin cleavage, and to stabilize prefusogenic conformations (11, 13, 51, 80–84). The glycosylation profile of virion S glycoproteins and several of these modified S glycoproteins has been characterized (30, 33, 36, 51, 70, 71). Our results agree with the overall predominance of complex carbohydrates of the SARS-CoV-2 S glycoprotein trimer seen in these previous studies. The glycosylation profile of our S glycoprotein preparation most closely resembles that of the S glycoproteins purified from SARS-CoV-2 virions propagated in Vero cells (52). However, compared with these and the other characterized trimers, the wild-type S glycoproteins purified in our study exhibited more glycan processing. The inclusion of a fucose-specific lectin in our purification scheme may have increased the representation of S glycoproteins that have passed through the Golgi, where complex carbohydrates are added (69). Soluble or virion-associated S glycoproteins passing through the Golgi may be processed less efficiently than our Golgi-enriched membrane-anchored S glycoproteins. The observed differences are particularly noteworthy for the Asn 234 glycan, which is predominantly of the high-mannose type in the soluble/modified S glycoproteins, but mostly processed in the wild-type S glycoprotein that we studied. Asn 234 in the S1 N-terminal domain is near the receptor-binding domain, and molecular dynamics simulations have suggested that the N234 glycan can modulate the conformational changes that the receptor-binding domain undergoes in the process of binding ACE2 (85). Moreover, changes in Asn 234 have been reported to affect virus sensitivity to several neutralizing antibodies (86). The particular types of glycans modifying Asn 234 might influence the binding and/or generation of neutralizing antibodies directed against nearby epitopes. Whatever its function, our results indicate that the N234 glycan is accessible on the unliganded wild-type S glycoprotein trimer for modification to complex carbohydrates. Specific down-selection of the N234 high-mannose glycans by AAL purification is possible but not likely, given the abundance of multiple fucose-containing glycans on all S glycoforms produced in the 293T-S cells.

O-linked glycans were detected on four glycopeptides in the purified S glycoprotein. Changing Thr 323, Thr 676 and Ser 1170 to those amino acid residues found in less common natural SARS-CoV-2 variants resulted in entry-competent S glycoproteins. However, the infectivity of the T323I mutant was more sensitive to freeze-thawing than that of viruses pseudotyped with the wild-type S glycoprotein. On the other hand, even though the S1170F change altered post-translational modification of the S2 glycoprotein, this mutant exhibited wild-type levels of infectivity and resistance to freeze-thawing. Alteration of Thr 676 or Ser 1170 did not significantly change the sensitivity of the pseudotyped viruses to neutralization by sACE2 or convalescent-phase sera.

Of the three naturally observed variants in S glycoprotein cysteine residues, a change in Cys 15 was compatible with an entry-competent S glycoprotein. This implies that the disulfide bond between Cys 15 and Cys 136 within the N-terminal domain is not absolutely required for folding and function of the SARS-CoV-2 S glycoprotein. However, we noted that the infectivity of the C15F mutant virus was compromised after freeze-thawing more than that of the wild-type virus. Apparently, some flexibility in the N-terminal domain can be tolerated in functional S glycoprotein trimers, although the ability of the virus to withstand environmental stress may be affected. Of note, some of the expressed S glycoproteins formed a disulfide bond between Cys 131 and Cys 136, and therefore lacked two of the canonical disulfide bonds (Cys 15-Cys 136 and Cys 131-Cys 166) in the N-terminal domain. Such S conformers, with presumably less stable N-terminal domains, might potentially contribute to viral pathogenesis or to evasion of the host immune response.

These studies should assist understanding of the nature and contribution of glycans on the wild-type SARS-CoV-2 S glycoprotein trimer and provide some insight into the impact of natural variation in sites that are glycosylated or disulfide bonded.

## EXPERIMENTAL PROCEDURES

### Reagents

Trizma^©^ hydrochloride, Trizma^©^ base, ammonium bicarbonate, urea, tris(2-carboxyethyl) phosphine hydrochloride (TCEP), iodoacetamide (IAM), ethanol, 4-vinylpyridine (4-VP), and glacial acetic acid were purchased from Sigma. Other reagents used in this study included optima LC/MS grade acetonitrile, water, formic acid (Fisher Scientific), sequencing grade trypsin (Promega), chymotrypsin (Promega), glycerol-free peptidyl-N-glycosidase F (PNGase F) (New England Biolabs), Endoglycosidase Hf (Endo Hf) (New England Biolabs), O-glycosidase (New England Biolabs), neuraminidase (New England Biolabs) and fetuin (New England Biolabs). All reagents and buffers were prepared with deionized water purified with a Millipore Direct-Q3 (Billerica, MA) water purification system.

### Plasmids

The wild-type and mutant SARS-CoV-2 S glycoproteins were expressed transiently by a pcDNA3.1(-) vector (Thermo Fisher Scientific) (29). The wild-type SARS-CoV-2 spike (S) gene sequence, which encodes an aspartic acid residue at position 614, was obtained from the National Center for Biological Information (NC_045512.20). The gene was modified to encode a Gly_3_ linker and His_6_ tag at the carboxyl terminus. The modified S gene was codon-optimized, synthesized by Integrated DNA Technologies, and cloned into the pcDNA3.1(-) vector. S mutants were made using Q5 High-Fidelity 2X Master Mix and KLD Enzyme Mix for site-directed mutagenesis, according to the manufacturer’s protocol (New England Biolabs), and One-Shot TOP10 Competent Cells.

Inducible expression of the wild-type SARS-CoV-2 S glycoprotein was achieved using a self-inactivating lentivirus vector comprising TRE.3g-SARS-CoV-2-Spike.6His.IRES6A.Puro.T2A.GFP (K5650) (29). Here, the codon-optimized *S* gene is under the control of a tetracycline response element (TRE) promoter and encodes the wild-type S glycoprotein with a carboxy-terminal 2xStrep tag. The internal ribosome entry site (IRES6A) allows expression of puro.T2A.EGFP, in which puromycin N-acetyltransferase and enhanced green fluorescent protein (eGFP) are produced by self-cleavage at the Thosea asigna 2A (T2A) sequence.

### Cell lines

The wild-type SARS-CoV-2 S glycoprotein, with Asp 614, was inducibly expressed in Lenti-x-293T human female kidney cells from Takara Bio (Catalog #: 632180). Lenti-x-293T cells were grown in DMEM with 10% FBS supplemented with L-glutamine and Pen-Strep.

Lenti-x-293T cells constitutively expressing the reverse tetracycline-responsive transcriptional activator (rtTA) (Lenti-x-293T-rtTa cells (D1317)) (29) were used as the parental cells for the 293T-S cell line. The 293T-S (D1483) cells inducibly expressing the wild-type SARS-CoV-2 S glycoprotein with a carboxy-terminal 2xStrep-Tag II sequence (29) were produced by transduction of Lenti-x-293T-rtTA cells with the K5650 recombinant lentivirus vector described above. The packaged K5650 lentivirus vector (60 µl volume) was incubated with 2×10^5^ Lenti-x-293T-rtTA cells in DMEM, tumbling at 37°C overnight. The cells were then transferred to a 6-well plate in 3 ml DMEM/10% FBS/Pen-Strep and subsequently selected with 10 µg/ml puromycin.

The GALE/GALK2 cells were obtained from Kerafast (73).

### Expression and processing of S glycoprotein variants

293T cells were transfected with plasmids expressing the wild-type and mutant SARS-CoV-2 S glycoproteins. On the day prior to transfection, 293T cells were seeded in 6-well plates at a density of 1 x 10^6^/well. Cells were transfected with 1 microgram of S-expressing plasmid, using Lipofectamine 3000 according to the manufacturer’s instructions. Two days after transfection, cells were lysed with lysis buffer (1x PBS, 1% NP-40 and 1x protease inhibitor cocktail) and the cell lysates analyzed by Western blotting. Samples were Western blotted with 1:2,000 dilutions of rabbit anti-SARS-Spike S1, mouse anti-SARS-Spike S1, rabbit anti-SARS-Spike S2 or a 1:5,000 dilution of mouse anti-β-actin as the primary antibodies. HRP-conjugated anti-rabbit or anti-mouse antibodies at a dilution of 1:5,000 were used as secondary antibodies in the Western blots. The adjusted integrated volumes of S, S1 and S2 bands from unsaturated Western blots were calculated using Fiji ImageJ. The values for the processing of mutant S glycoproteins were calculated and normalized to the values for the wild-type S glycoprotein (WT) as follows:

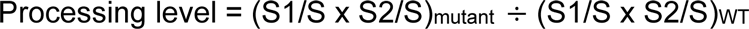

For production of VLPs, the SARS-CoV-2 S glycoprotein was coexpressed with M, E and N proteins, individually or in combination. One day before transfection, 293T cells were seeded into 10-cm dishes at a density of 5.5 x 10^6^/dish. The next day, the cells were transfected with 3 micrograms of each expressor plasmid or with an empty vector plasmid to keep the total amount of DNA transfected at 12 micrograms. Two days after transfection, cell lysates were prepared as described above. Cell supernatants were cleared at 900 x g for 15 minutes, followed by centrifugation at 100,000 x g for one hour at 4°C. Pellets were washed with 800 µl of 1x PBS and then resuspended in 90 µl lysis buffer for 5 minutes on ice. In some cases, cell lysates and pellets prepared from cell supernatants were treated with PNGase F, Endoglycosidase Hf (Endo Hf), or O-glycosidase + neuraminidase (New England Biolabs), according to the manufacturer’s instructions. Samples were analyzed by Western blotting as described above.

### S1 shedding from S glycoprotein-expressing cells

293T cells were transfected with pcDNA3.1(-) plasmids expressing the wild-type and mutant SARS-CoV-2 S glycoproteins, using Lipofectamine 3000 according to the manufacturer’s protocol. Cell supernatants were collected, cleared by centrifugation at 1800 x g for 10 minutes and incubated with a 1:100 dilution of NYP01 convalescent serum and Protein A-agarose beads for 1-2 hours at room temperature. Beads were washed three times and samples were Western blotted with a mouse anti-S1 antibody. Band intensity was determined as described above. The subunit association index of each mutant was calculated as follows:

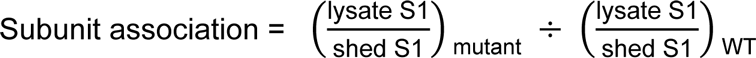

### Purification of the S glycoproteins

To express the SARS-CoV-2 S glycoprotein for purification, 293T-S cells were induced with 1 µg/ml doxycycline for two days. The cells were resuspended in 1x PBS and spun at 4500 x g for 15 minutes at 4°C. Cell pellets were collected and lysed by incubating in lysis buffer (20 mM Tris HCl (pH 8.0), 150 mM NaCl, 1% Cymal-5, 1x protease inhibitor cocktail (Roche)) on ice for 10 minutes. Cell lysates were spun at 10,000 x g for 20 minutes at 4°C, and the supernatant was incubated with Strep-Tactin XT Superflow resin (IBA # 2-4030-010) by rocking end over end at room temperature for 1.5 hours in a 50-ml conical tube. After incubation, the supernatant-resin suspension was applied to a Biorad column allowing flowthrough by gravity, followed by washing with 20 bed volumes of washing buffer (IBA # 2-1003-100 containing 0.5% Cymal-5), and elution with 10 bed volumes of elution buffer (IBA # 2-1042-025 containing 0.5% Cymal-5 and 1x protease inhibitor cocktail). For the second step of purification, the eluate was incubated with AAL-agarose resin (Vector Laboratories # AL-1393-2) at room temperature for 1 hour in a 10-ml conical tube. The eluate-AAL resin suspension was applied to a Biorad eco-column for gravity flowthrough. The column was washed with 20 bed volumes of washing buffer (20 mM Tris HCl (pH 8.0), 150 mM NaCl, 0.5% Cymal-5, 1x protease inhibitor cocktail (Roche)), after which the sample was eluted with 10 bed volumes of elution buffer (9 parts elution buffer (Vector Laboratories # ES-3100-100), 0.5 parts 1 M Tris HCl (pH 8.0), 0.5 parts 10% Cymal-5). The eluate was buffer exchanged by ultrafiltration three times to remove fucose; this was accomplished using a 15-ml ultrafiltration tube (Thermo Fisher Scientific # UFC903024) at 4000 x g at room temperature with a buffer consisting of 20 mM Tris HCl (pH 8.0), 150 mM NaCl and 0.5% Cymal-5.

### Proteolytic digestion of SARS-CoV-2 spike glycoproteins for glycosylation analysis

The purified SARS-CoV-2 S glycoprotein samples (30 µg) at a concentration of ∼0.03 mg/mL were denatured with 7 M urea in 100 mM Tris buffer (pH 8.5), reduced at room temperature for one hour with TCEP (5 mM), and alkylated with 20 mM IAM at room temperature for another hour in the dark. The reduced and alkylated samples were buffer exchanged with 50 mM ammonium bicarbonate (pH 8) using a 50-kDa MWCO filter (Millipore) prior to trypsin digestion. The resulting buffer-exchanged sample was aliquoted into two portions – one digested with trypsin and the other with chymotrypsin. All protease digestions were performed according to manufacturer’s suggested protocols: digestion with trypsin was performed with a 30:1 protein:enzyme ratio at 37°C for 18 hours; chymotrypsin digestion was performed with a 20:1 protein:enzyme ratio at 30°C for 10 hours; the combination of both proteases (a mixture of trypsin and chymotrypsin) was performed using the same protein:enzyme ratio as that used for single enzyme digestion and was incubated overnight at 37°C. Ten microliter aliquots from each digest were treated with PNGase F and incubated at 37°C. The digests were either directly analyzed or stored at −20°C until further analysis.

### Disulfide bond analysis of SARS-CoV-2 spike glycoprotein

The disulfide bond profiles of the SARS-CoV-2 spike glycoproteins were determined as described previously (87, 88). Briefly, a sample containing 10 μg of the spike glycoprotein was buffer exchanged with 50 mM ammonium citrate buffer (pH 6.5) using a 50-kDa MWCO filter (Millipore). The resulting buffer-exchanged sample was alkylated with a 20-fold molar excess of 4-vinylpyridine in the dark for one hour at room temperature to cap free cysteine residues. The alkylated sample was subsequently deglycosylated with 500 U of PNGase F for one week at 37°C. The deglycosylated and alkylated sample was digested overnight with trypsin (protein to enzyme ratio of 30:1 at 37°C. A 20-µL aliquot from the tryptic digest was further treated with 1 µg chymotrypsin and was incubated for eight hours at 37°C. The digests were either directly analyzed or stored at −20°C until further analysis.

### Chromatography and mass spectrometry

High-resolution LC/MS experiments were performed using an Orbitrap Fusion Lumos Tribrid (Thermo Scientific) mass spectrometer equipped with ETD that is coupled to an Acquity UPLC M-Class system (Waters). Mobile phases consisted of solvent A: 99.9% deionized H_2_O + 0.1% formic acid and solvent B: 99.9 % CH_3_CN + 0.1% formic acid. Three microliters of the sample was injected onto a C18 PepMap™ 300 column (300 µm i.d. x 15 cm, 300 Å, Thermo Fisher Scientific) at a flow rate of 3 µL/min. The following CH_3_CN/H_2_O multistep gradient was used: 3% B for 3 min, followed by a linear increase to 45% B in 50 min then a linear increase to 90% B in 15 min. The column was held at 90% B for 10 minutes before re-equilibration. All mass spectrometric analysis was performed in the positive ion mode using data-dependent acquisition with the instrument set to run in 3-sec cycles for the survey and two consecutive MS/MS scans with CID and EThcD or ETciD. The full MS survey scans were acquired in the Orbitrap in the mass range 400-1800 *m/z* at a resolution of 120000 at *m/z* 200 with an AGC target of 4 x10^5^. Following a survey scan, MS/MS scans were performed on the most intense ions with charge states ranging from 2-6 and with intensity greater than 5000. CID was carried out at with a collision energy of 30% while ETD was performed using the calibrated charge-dependent reaction time. Resulting fragments were detected using rapid scan rate in the ion trap.

### Glycopeptide identification and disulfide bond analysis

Glycopeptide compositional analysis was performed as described previously (89, 90). Briefly, compositional analysis of glycopeptides was carried out by first identifying the peptide portion from tandem MS data. Once the peptide portion was determined, plausible glycopeptide compositions were obtained using the high-resolution MS data and GlycoPep DB (91). The putative glycopeptide composition was confirmed manually from CID and ETD data.

Disulfide bond patterns of SARS CoV-2 spike glycoproteins were determined by mapping the disulfide-linked peptides. Data analysis was performed using the Mascot (v2.5.1) search engine (92) for peptides containing free cysteine residues, and disulfide bond patterns were analyzed manually, as described previously (87, 88).

### VSV pseudotyped by S glycoproteins

VSV was pseudotyped with S glycoproteins expressed stably in 293T-S cells or transiently in 293T cells. 293T-S cells in 6-well plates were induced with 1 µg/ml doxycycline or, as a control, incubated in standard medium without doxycycline. For transient expression, subconfluent 293T cells in a T75 flask were transfected with 15 µg of the SARS-CoV-2 S expressor plasmid using 60 µl of 1 mg/ml polyethylenimine (PEI). Twenty-four hours later, cells were infected at a multiplicity of infection of 3-5 for 2 hours at 37°C with rVSV-ΔG pseudovirus complemented in *trans* with the G glycoprotein and bearing a luciferase gene (Kerafast). Cells were then washed 6 times with DMEM + 10% FBS and returned to culture. Cell supernatants containing S-pseudotyped VSV were harvested 24 hours later, clarified by low-speed centrifugation (900 x g for 10 min), and either characterized immediately or stored at −80°C for later analysis.

### Syncytium formation assay

293T-S cells in 6-well plates were cotransfected with 1 µg each of an eGFP-expressing plasmid and a plasmid expressing hACE2, and then incubated in either standard (control) medium or medium containing 1 µg/ml doxycycline. Twenty-four hours after transfection, cells were imaged using a fluorescence microscope with a green light filter. In parallel, cell lysates were collected for Western blotting, as described above.

### Virus infectivity

VSV-ΔG vectors pseudotyped with SARS-CoV-2 S gp variants were produced as described above. The recombinant viruses were incubated with 293T-ACE2 cells, and 24 hours later, luciferase activity in the cells was measured.

### Virus neutralization by sACE2 and sera

Neutralization assays were performed by adding 200-300 TCID_50_ of rVSV-ΔG pseudotyped with SARS-CoV-2 S glycoprotein variants into serial dilutions of sACE2 and sera. The mixture was dispensed onto a 96-well plate in triplicate and incubated for 1 h at 37°C. Approximately 4 x 10^4^ 293T-ACE2 cells were then added to each well, and the cultures were maintained for an additional 24 h at 37°C before luciferase activity was measured. Neutralization activity was calculated from the reduction in luciferase activity compared to controls, using GraphPad Prism 8 (GraphPad Software Inc.).

## DATA AVAILABILITY

All the data supporting the conclusions are contained in the manuscript.

## ACKNOWLEDGMENTS

We thank Ms. Elizabeth Carpelan for preparation of the manuscript. We thank Lihong Liu and David Ho (Columbia University), Peihui Wang (Shandong University) and Yuan Liu (Cornell University) for reagents.

## FUNDING AND ADDITIONAL INFORMATION

This study was supported by the University of Alabama at Birmingham Center for AIDS Research (NIH P30 AI27767), by grants from the National Institutes of Health (AI125093 to H.D., J.C.K. and J.S. and R35 GM130354 to H.D.), and by a gift to J.S. from the late William F. McCarty-Cooper. The content is solely the responsibility of the authors and does not necessarily represent the official views of the National Institutes of Health.

